# A Pilot Evaluation of Open-Weight Large Language Models for Screening RNA-seq Metadata in Public Databases

**DOI:** 10.64898/2026.02.16.706241

**Authors:** Mitsuo Shintani, Daniel Andrade, Hidemasa Bono

**Author notes:** Correspondence: Hidemasa Bono.

## Abstract

Although the Gene Expression Omnibus and other public repositories are expanding rapidly, curation across these databases has not kept pace. Data reuse is often hindered by unstandardized metadata comprising unstructured text. To address this, we developed a workflow that combines retrieval via application programming interfaces with semantic filtering using large language models (LLMs) to support metadata screening as an initial step in broader curation workflows. As a focused pilot evaluation, we benchmarked multiple LLMs using metadata from 150 candidate *Arabidopsis* RNA-seq projects to classify projects containing exogenous ABA-treated samples and matched untreated controls. Simple keyword searches yielded many false positives (F1=0.59); classification using LLMs significantly improved performance. Several open-weight models achieved near-perfect classification performance in this defined task (F1>0.98), comparable to that of closed models. We also found that, for some high-performing models, self-reported confidence scores may help identify high-confidence cases that can be prioritized for automated processing. These results suggest that open-weight LLMs can support scalable metadata screening in local environments as an initial step in broader curation workflows, providing a foundation for accelerating public dataset reuse.

## INTRODUCTION

In recent years, scientific data has grown rapidly as experimental, observational, and computational technologies have advanced. However, metadata standardization and curation practices that enable data reuse have not kept pace with this growth ^1^. The rapid development of next-generation sequencing technology has transformed life science research. RNA sequencing (RNA-seq) has enabled the comprehensive quantitative measurement of all RNA transcripts (the transcriptome) present in a specific cell type or tissue and has become a major approach for studying gene expression^2^. With the spread of this technology, large amounts of RNA-seq data obtained from many species, tissues, and experimental conditions are being deposited daily in public databases, such as the NCBI Sequence Read Archive (SRA) and the Gene Expression Omnibus (GEO) ^3,4^. These public datasets are important resources for data reuse and reanalysis, including meta-analysis ^5,6^. However, effective reuse requires accurate identification of datasets that match specific biological and experimental criteria, which remains difficult when metadata are unstandardized and described in heterogeneous natural language. In practice, this difficulty is not limited to retrieving records that contain relevant keywords. Key experimental information such as organism, assay type, treatment, control availability, tissue, genotype, and sample correspondence is often distributed across project- and sample-level descriptions and is not always recorded in standardized fields.

Meanwhile, many efforts have been made to lower the barriers to reusing public data. For GEO, GEOmetaDB has been proposed as a local database and query layer that enables more flexible and efficient metadata searches than standard web interfaces ^7^. For SRA, programmable tools such as PysRADB support the large-scale retrieval of study and sample metadata from the command line ^8^. PubTator assists in biocuration by automatically extracting and normalizing major entities from PubMed articles and providing them as pre-annotations ^9^. More recently, PubTator 3.0 enables a semantic and relational search over biomedical literature using artificial intelligence (AI) techniques, and it further demonstrates that integrating PubTator application programming interfaces (APIs) with a large language model (LLM) can support AI-assisted literature exploration and improve the factuality and verifiability of evidence-based responses ^10^. These tools are useful for metadata access, retrieval, and organization. However, an additional bottleneck remains after retrieval: determining whether each retrieved project satisfies user-defined eligibility criteria for downstream reanalysis.

Keyword-based searches may retrieve projects that mention a treatment term in a description, sample label, or related context, even when the project does not contain the intended treated samples and matched controls. Thus, improving search and access alone is often insufficient to facilitate smooth reanalysis in practice, and it is also critical to streamline the process of accurately selecting datasets that match analysis goals. Because much of the metadata essential for analysis is recorded in inconsistent natural language across project- and sample-level descriptions, this downstream judgment requires semantic interpretation and has largely remained dependent on manual labor. This extensive, repetitive workload is time- and labor-intensive, and acts as a significant barrier to scaling and accelerating data-driven research (Figure 1a).

**Figure 1:**
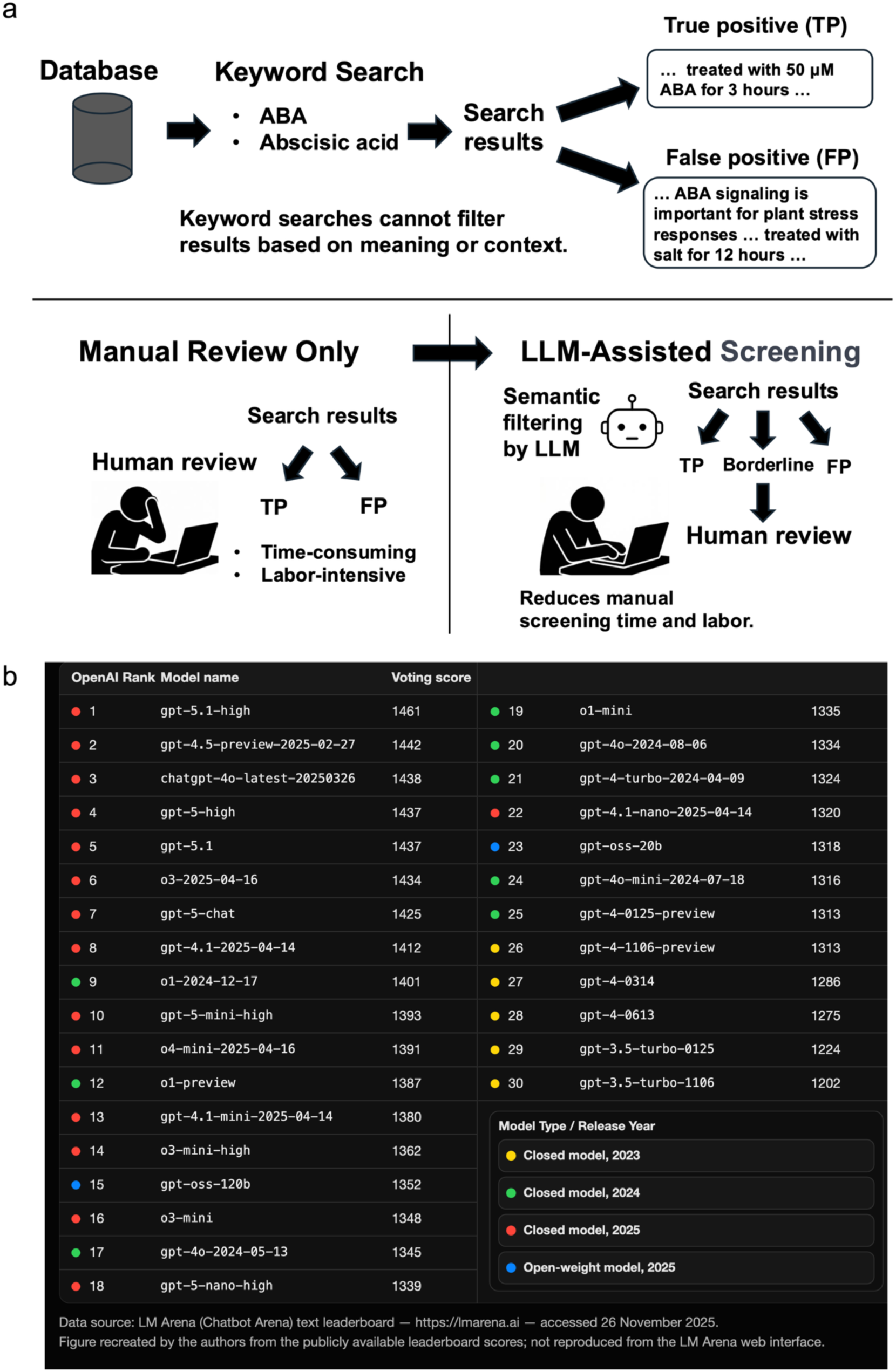
Motivation for LLM-assisted metadata screening and the evolution of model performance. (a) Keyword searches cannot filter based on meaning or context, which leads to false positives and a heavy workload for manual reviews. LLM-assisted screening enables semantic filtering of search results, reducing the time and labor required for manual review. (b) Performance scores from the LM Arena text leaderboard for OpenAI’s closed and open-weight models released between November 2023 and November 2025, with newer open-weight models tending to match or exceed the performance of earlier closed models. Figure 1b was recreated by the authors using score data obtained from the LM Arena text leaderboard. Data were sourced from LM Arena and accessed on November 26, 2025.

LLMs learn from large text corpora and can capture complex patterns in natural languages, enabling advanced operations such as context-aware text generation, classification, and summarization ^11^.

LLMs do not understand meaning in the same way as humans do, but their ability to recognize statistical patterns allows them to process information written in natural language. Using this ability, it may be possible to automate and accelerate information extraction and classification from unstructured metadata, which has previously relied on manual work. A key turning point was the release of GPT-4 in 2023, which marked the beginning of a clear expansion in the practical capabilities of LLMs. ^12^.

Recent studies have reported LLM-based approaches for extracting and standardizing biological metadata from unstructured text. For example, in a task that extracts biological terms (e.g., cell line names) from BioSample descriptions, LLM-assisted methods have been shown to improve extraction performance compared with traditional rule-based methods ^13^. Other studies explored LLM-based assignment of ontology terms for biological sample annotation, suggesting that under appropriate constraints, generative models can support curation workflows that consider ontologies ^14^. Taken together, these studies suggest that the use of LLMs to reduce the manual workload in metadata curation is a realistic possibility. At the same time, factors such as output accuracy, outputs that follow user-defined standards and formats, and application to large-scale data remain key challenges for practical deployment.

Open-weight models are attractive for research workflows because the model version can be fixed and executed in a local environment, which may improve reproducibility and reduce dependence on provider-managed APIs. Recent progress in quantization and local inference frameworks has also made it increasingly feasible to run mid-scale models on local workstations. In this study, we therefore evaluated whether locally executable open-weight models can achieve sufficient accuracy for metadata screening tasks.

Open-weight models have also improved rapidly. In some cases, within months, they can achieve a performance comparable to that of earlier state-of-the-art closed models. LM Arena (formerly Chatbot Arena) evaluates model performance by running the same tasks with two anonymized models on a website and collecting user votes in an A/B test setting ^15,16^. Figure 1b was recreated by the authors using score data obtained from the LM Arena text leaderboard. Data were sourced from LM Arena and accessed on November 26, 2025. These scores were taken from the LM Arena text leaderboard as November 26, 2025. The models shown as blue points, gpt-oss-20B and gpt-oss-120B, are open-weight models released by OpenAI in August 2025 ^17^. These models show performances that are comparable to or higher than those of the closed models that were released and used during 2023–2024 (shown as yellow or green points). This suggests that the open-weight models available in 2025 already have a performance comparable to past closed models and that it may be possible to run tasks with an accuracy that meets or exceeds that of past closed models.

In this study, we tested whether part of the metadata screening work for public datasets can be automated using multiple language models, including the open-weight model gpt-oss-120B, which can run in a local environment. We focused on reducing false positives remaining after keyword-based retrieval by applying LLM-based semantic classification to project- and sample-level metadata.

We then evaluated how accurately LLMs could perform such tasks and how much they could contribute to automation. We also used closed models released in the past two years via APIs and compared their accuracy with that of open-weight models to assess the extent to which the open-weight model performance has advanced.

## RESULTS

### Development of an LLM-assisted metadata screening workflow for public RNA-seq data

In this study, we present a workflow that supports initial metadata screening for public RNA-seq projects by combining API-based metadata retrieval with LLM-based semantic filtering. Figure 2 summarizes the conceptual phases of the LLM-assisted metadata screening and evaluation workflow, visually illustrating the three phases described in the text: broadly collecting candidate studies through keyword search, automatically determining experimental relevance and sample composition using an LLM, and finally comparing the classification accuracy and processing speed of each model.

**Figure 2:**
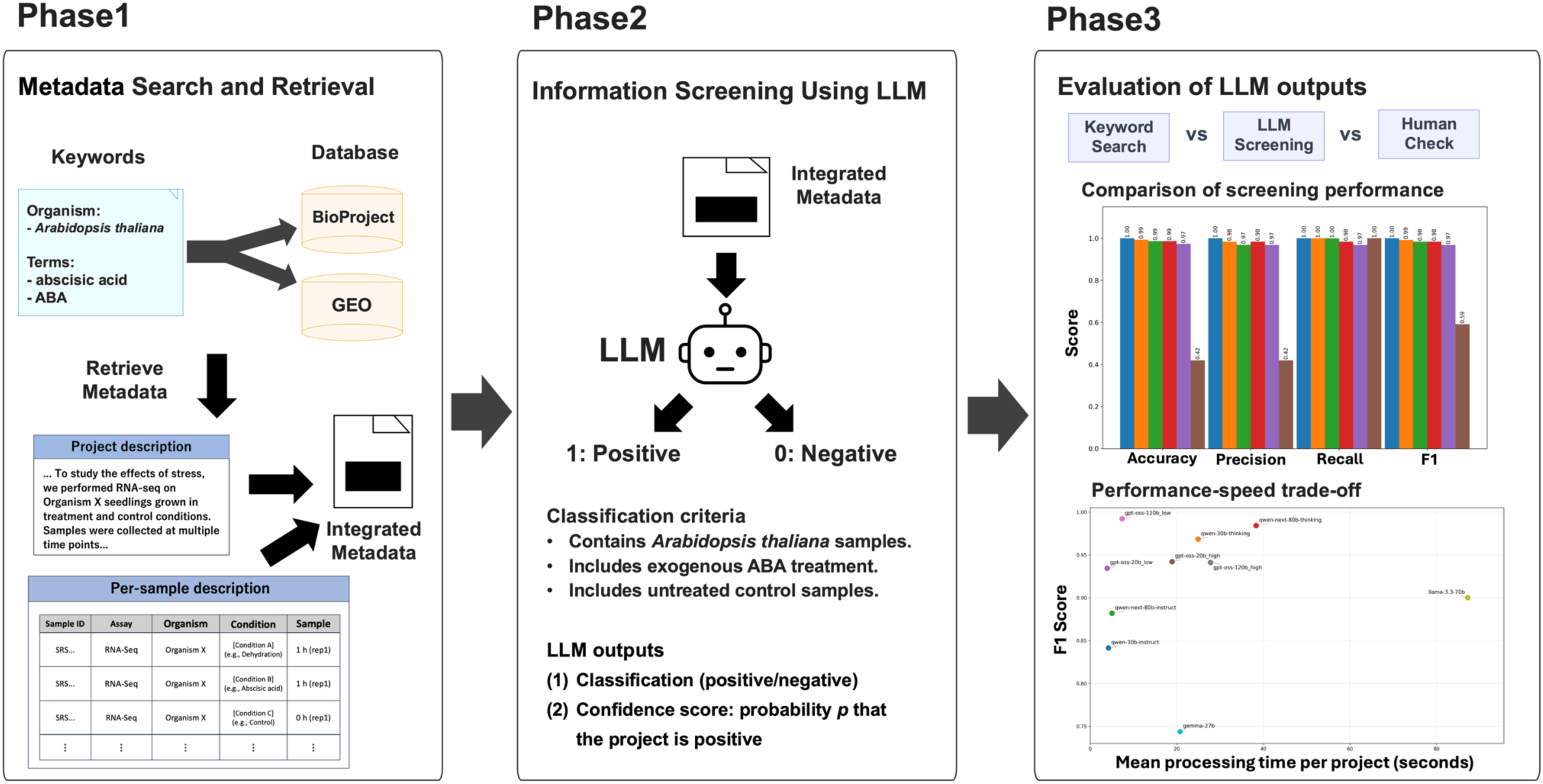
Conceptual phases of the LLM-assisted metadata screening and evaluation workflow. The pipeline programmatically retrieves project- and sample-level metadata via public APIs and consolidates them into a structured text input. An LLM then performs semantic filtering to identify projects containing *Arabidopsis thaliana* RNA-seq samples with exogenous ABA treatment and matched untreated controls, while outputting a confidence score (*p*), and model performance/runtime are benchmarked across local open-weight and API-based closed models. These phases are distinct from the script execution steps described in the GitHub 04_workflow.

Although these phases broadly correspond to the workflow provided on GitHub, they are intended to represent the conceptual organization of the study rather than the exact script execution sequence.

### LLM-based metadata classification results

Next, we describe the results of metadata classification using LLMs. Using a benchmark dataset of 150 projects, we ran the same metadata classification task with multiple LLMs and evaluated their output accuracy. For evaluation, we used four commonly used classification metrics: accuracy, precision, recall, and F1 score; the results are shown in Table 1. Each metric takes a value between 0 and 1, with higher values indicating better performance. Accuracy represents the fraction of all samples that were correctly classified, precision indicates the suppression of false positives (FPs), recall indicates the suppression of false negatives (FNs), and the F1 score evaluates the balance between precision and recall.

**Table 1:** Comparison of model accuracy on the classification task using two prompts. Performance metrics of each model under prompt 1 and prompt 2, presented together with a baseline using keyword search only.

| Model | Prompt version | Accuracy | Precision | Recall | F1 score |
| --- | --- | --- | --- | --- | --- |
| gemini-2.5-pro | 1 | 0.960 | 0.913 | 1.000 | 0.955 |
| gemini-2.5-pro | 2 | 1.000 | 1.000 | 1.000 | 1.000 |
| gemma-3-27b-it-qat | 1 | 0.720 | 0.602 | 0.984 | 0.747 |
| gemma-3-27b-it-qat | 2 | 0.720 | 0.604 | 0.968 | 0.744 |
| gpt-3.5-turbo-0125 | 1 | 0.560 | 0.488 | 0.952 | 0.645 |
| gpt-3.5-turbo-0125 | 2 | 0.547 | 0.479 | 0.921 | 0.630 |
| gpt-4o-2024-11-20 | 1 | 0.640 | 0.539 | 1.000 | 0.700 |
| gpt-4o-2024-11-20 | 2 | 0.847 | 0.733 | 1.000 | 0.846 |
| gpt-4o-mini-2024-07-18 | 1 | 0.500 | 0.456 | 0.984 | 0.623 |
| gpt-4o-mini-2024-07-18 | 2 | 0.553 | 0.485 | 1.000 | 0.653 |
| gpt-5.1-2025-11-13 | 1 | 0.913 | 0.829 | 1.000 | 0.907 |
| gpt-5.1-2025-11-13 | 2 | 0.987 | 0.969 | 1.000 | 0.984 |
| gpt-oss-safeguard-120b | 1 | 0.967 | 0.927 | 1.000 | 0.962 |
| gpt-oss-safeguard-120b | 2 | 0.973 | 0.984 | 0.952 | 0.968 |
| gpt-oss-safeguard-20b | 1 | 0.947 | 0.899 | 0.984 | 0.939 |
| gpt-oss-safeguard-20b | 2 | 0.953 | 0.983 | 0.905 | 0.942 |
| meta/llama-3.3-70b | 1 | 0.673 | 0.563 | 1.000 | 0.720 |
| meta/llama-3.3-70b | 2 | 0.907 | 0.818 | 1.000 | 0.900 |
| openai/gpt-oss-120b_high | 1 | 0.973 | 0.954 | 0.984 | 0.969 |
| openai/gpt-oss-120b_high | 2 | 0.953 | 1.000 | 0.889 | 0.941 |
| openai/gpt-oss-120b_low | 1 | 0.967 | 0.953 | 0.968 | 0.961 |
| openai/gpt-oss-120b_low | 2 | 0.993 | 0.984 | 1.000 | 0.992 |
| openai/gpt-oss-20b_high | 1 | 0.940 | 0.909 | 0.952 | 0.930 |
| openai/gpt-oss-20b_high | 2 | 0.953 | 0.983 | 0.905 | 0.942 |
| openai/gpt-oss-20b_low | 1 | 0.933 | 0.884 | 0.968 | 0.924 |
| openai/gpt-oss-20b_low | 2 | 0.947 | 0.966 | 0.905 | 0.934 |
| **qwen3-30b-a3b-instruct-2507** | 1 | 0.660 | 0.553 | 1.000 | 0.712 |
| **qwen3-30b-a3b-instruct-2507** | 2 | 0.847 | 0.744 | 0.968 | 0.841 |
| **qwen3-30b-a3b-thinking-2507** | 1 | 0.940 | 0.886 | 0.984 | 0.932 |
| **qwen3-30b-a3b-thinking-2507** | 2 | 0.973 | 0.968 | 0.968 | 0.968 |
| **qwen3-next-80b-a3b-instruct** | 1 | 0.647 | 0.543 | 1.000 | 0.704 |
| **qwen3-next-80b-a3b-instruct** | 2 | 0.900 | 0.875 | 0.889 | 0.882 |
| **qwen3-next-80b-a3b-thinking** | 1 | 0.900 | 0.808 | 1.000 | 0.894 |
| **qwen3-next-80b-a3b-thinking** | 2 | 0.987 | 0.984 | 0.984 | 0.984 |
| **Keyword Search Only (Baseline)** | - | 0.420 | 0.420 | 1.000 | 0.592 |

To compare the performance among LLMs, we also evaluated against a method using only keyword search. Specifically, we assumed the model “Keyword Search Only (Baseline)” that classifies all projects returned by keyword search as positive, and we computed the same metrics and reported them alongside the LLM results. Under this baseline, the recall value was 1.00 by definition within the retrieved candidate pool, because all projects returned by the keyword search were treated as positive. This value does not indicate that the keyword search retrieved all positive projects from the entire database. By comparing against this baseline, we can assess how much semantic classification via LLM reduces the FPs arising from keyword searches alone. We compared a prompt that lists only the minimal criteria to avoid missing positive projects (prompt 1) with a prompt that adds detailed criteria to suppress false positives (prompt 2). Table 1 shows the classification performance of each model using prompts 1 and 2, as well as the baseline values. The baseline condition using keyword search only yielded an accuracy of 0.42, precision of 0.42, recall of 1.00, and F1 of 0.59, showing that while recall was maximal, precision was low, and many FPs were included. All LLMs substantially exceeded these metrics. The highest performance was achieved by gemini-2.5-pro with prompt 2, which attained a value of 1 for all metrics and produced a classification that perfectly matched the ground-truth labels. These results confirm that compared with extracting candidates via keyword search alone, performing LLM-based classification on candidates obtained from keyword searches markedly improves the overall performance. In other words, LLM-based semantic classification can effectively remove FPs produced by keyword searches and increase the extraction accuracy of projects that meet the specified criteria.

Meanwhile, the open-weight models released in 2025, such as gpt-oss-120b and gpt-oss-safeguard-120b, achieved higher classification performance than the model released in 2023, gpt-3.5-turbo-0125. Notably, these open-weight models also outperformed the models released in 2024, gpt-4o-2024-11-20 and gpt-4o-mini-2024-07-18. Furthermore, these open-weight models showed metric values comparable to or slightly lower than high-performance closed models such as the 2025 releases of gemini-2.5-pro and gpt-5.1-2025-11-13. This indicates that the level of task accuracy achievable using open-weight models has steadily improved in recent years. Consequently, these results suggest that currently available open-weight models may enable the execution of tasks such as metadata classification with high accuracy without necessarily relying on the latest closed models.

We also computed the ratio of the evaluation metrics obtained with prompt 2 to those obtained with prompt 1 (P2/P1) and the results are presented in Table 2. A ratio greater than 1.0 indicates that prompt 2 yielded higher scores, whereas a ratio less than 1.0 indicates that prompt 1 yielded higher scores.

**Table 2:** Ratios of model accuracy on the classification task using two prompts. To quantify performance changes due to the prompts, the ratio of output accuracy under prompt 2 to that under prompt 1 (P2/P1) is reported.

| Model | Accuracy (P2/P1) | Precision (P2/P1) | Recall (P2/P1) | F1 (P2/P1) |
| --- | --- | --- | --- | --- |
| gemini-2.5-pro | 1.04 | 1.10 | 1.00 | 1.05 |
| gemma-3-27b-it-qat | 1.00 | 1.00 | 0.98 | 1.00 |
| gpt-3.5-turbo-0125 | 0.98 | 0.98 | 0.97 | 0.98 |
| gpt-4o-2024-11-20 | 1.32 | 1.36 | 1.00 | 1.21 |
| gpt-4o-mini-2024-07-18 | 1.11 | 1.06 | 1.02 | 1.05 |
| gpt-5.1-2025-11-13 | 1.08 | 1.17 | 1.00 | 1.09 |
| gpt-oss-safeguard-120b | 1.01 | 1.06 | 0.95 | 1.01 |
| gpt-oss-safeguard-20b | 1.01 | 1.09 | 0.92 | 1.00 |
| meta/llama-3.3-70b | 1.35 | 1.45 | 1.00 | 1.25 |
| openai/gpt-oss-120b_high | 0.98 | 1.05 | 0.90 | 0.97 |
| openai/gpt-oss-120b_low | 1.03 | 1.03 | 1.03 | 1.03 |
| openai/gpt-oss-20b_high | 1.01 | 1.08 | 0.95 | 1.01 |
| openai/gpt-oss-20b_low | 1.01 | 1.09 | 0.93 | 1.01 |
| qwen3-30b-a3b-instruct-2507 | 1.28 | 1.35 | 0.97 | 1.18 |
| qwen3-30b-a3b-thinking-2507 | 1.04 | 1.09 | 0.98 | 1.04 |
| qwen3-next-80b-a3b-instruct | 1.39 | 1.61 | 0.89 | 1.25 |
| qwen3-next-80b-a3b-thinking | 1.10 | 1.22 | 0.98 | 1.10 |

P2/P1 represents the ratio obtained by dividing each metric value for prompt 2 by the corresponding value for prompt 1.

We next examined how changing the strictness of the classification criteria in the prompt affected the precision-recall trade-off. This comparison was not intended as an exhaustive prompt-engineering optimization, but as a limited analysis of whether a more stringent prompt could reduce false positives arising from ambiguous or incomplete public metadata. Prompt 1 listed the minimum positive criteria and was designed to avoid missing potentially relevant projects, whereas prompt 2 added stricter rules requiring explicit evidence of exogenous ABA treatment and corresponding untreated controls within the same project. Prompt 2 also encouraged conservative classification when the available project-level or sample-level metadata did not provide sufficient evidence. Comparing prompts 1 and 2, prompt 2 tended to exclude FPs more accurately and improve precision in many models; however, it was accompanied by a decrease in recall in some models, such as openai/gpt-oss-120b_high, indicating a tendency toward more cautious classification. In gemini-2.5-pro, gpt-5.1-2025-11-13, and openai/gpt-oss-120b_low, precision improved while recall was maintained, resulting in higher F1 scores. These results show that prompt strictness can shift the precision-recall trade-off, but the magnitude and direction of this effect depend on the model. Therefore, prompts designed to handle metadata ambiguity or missingness should be validated for each target task and model rather than assumed to generalize universally.

Figure 3 shows the relationship between precision and recall for each model. The y-axis represents precision, the x-axis represents recall, and the F1 score is defined as the harmonic mean of the two values. Each point in the figure represents one model: points higher in the plot indicate higher precision and points farther to the right indicate higher recall. Therefore, models located closer to the upper-right corner indicated a better balance between precision and recall and achieved higher F1 scores. The dashed lines represent the iso-F1 curves for F1 = 0.85, 0.90, 0.925, and 0.95, enabling a visual comparison of the relative performance in suppressing FPs and FNs.

**Figure 3:**
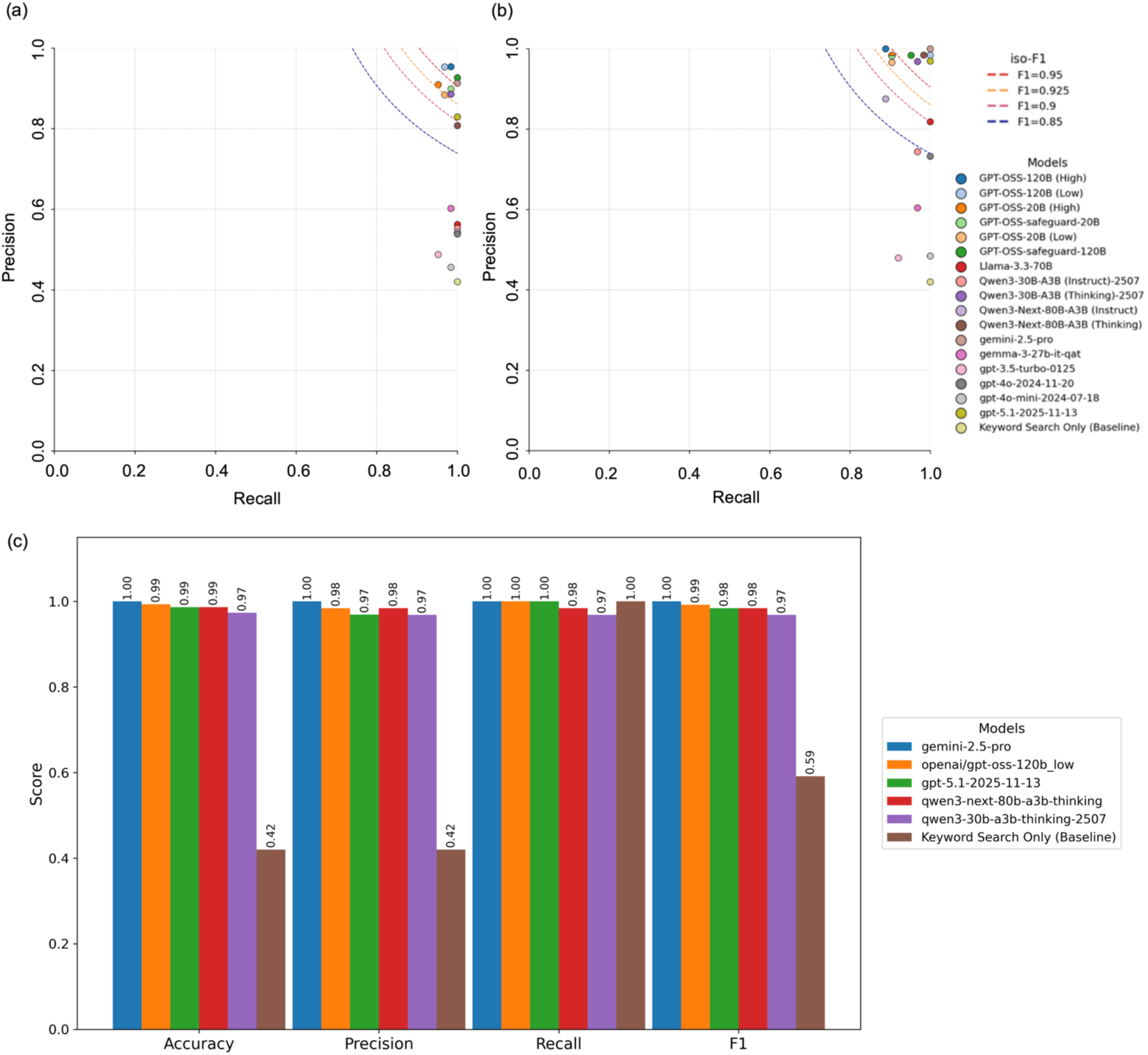
Precision–recall trade-offs for project-level screening across models and prompts. (a) Precision–recall plot for all evaluated models under prompt 1. Iso-F1 curves are shown for visual comparison. The plot illustrates how the precision–recall trade-off shifts depending on the model. (b) Precision–recall plot for all evaluated models under prompt 2. (c) Side-by-side comparison of accuracy, precision, recall, and F1 score for top-performing models. Candidate models that exceed high iso-F1 regions are highlighted.

When using prompt 1, the recall value was close to 1 for many models, whereas the precision values were often approximately 0.9. In contrast, when using prompt 2, precision increased to higher values, even for the same models, whereas the recall tended to decrease slightly. However, this trend was not uniform across the models. In Figure 3b, some models showed a large upward shift (precision improvement), whereas others showed a relatively large leftward shift (recall decrease). In particular, near the upper-right region around the iso-F1 curve of F1 ≥ 0.95 are gemini-2.5-pro (prompt 2), gpt-5.1-2025-11-13, openai/gpt-oss-120b_low, qwen3-next-80b-a3b-thinking, and qwen3-30b-a3b-thinking, indicating a group of models that can suppress both FPs and FNs at a high level. Overall, Figure 3 shows that prompt design can effectively move the precision-recall trade-off, while also showing that the manifestation of this effect is model-dependent.

Differences in performance were also observed within the same family of models. For example, in qwen3-30b and qwen3-next-80b, the thinking versions consistently showed higher F1 scores than the instruct versions. Specifically, for qwen3-30b, the F1 score was 0.712 (instruct) vs. 0.932 (thinking) with prompt 1, and 0.841 (instruct) vs. 0.968 (thinking) with prompt 2. Similarly, for qwen3-next-80b, the F1 score was 0.704 (instruct) vs. 0.894 (thinking) with prompt 1 and 0.882 (instruct) vs. 0.984 (thinking) with prompt 2, with the thinking versions outperforming the instruct versions. As shown in Figure 3a and 3b, the thinking versions are generally located closer to the upper right of the graph (high precision and high recall) than the instruct versions, suggesting that even within the same series, configurations that use reasoning processes more strongly showed improved classification performance (especially higher F1 score). Figure 3c presents a comparison of the four metrics for the models that achieved high F1 scores.

In summary, these results indicate that the LLM-based approach can achieve a precision-recall trade-off that is substantially superior to the keyword-search-only baseline. In particular, the group of models located in the upper-right region exceeding the iso-F1 curve of 0.95 are promising candidates for use in curation tasks similar to those in this study.

### Analysis of LLMs’ self-reported positive label probabilities

In addition to the binary labels, each model was asked to self-report the probability of a project being positive with a value between zero and one. Supplementary Figure 1 shows the distribution of the self-reported positive probabilities for each model (prompt 2).

Projects with probabilities near 0.5 were considered ambiguous inputs for the model, indicating that the decision between positive and negative was unclear. In this study, high-confidence predictions were predefined as p < 0.25 or p > 0.75, whereas predictions with 0.25 ≤ p ≤ 0.75 were treated as low-confidence or intermediate predictions and excluded from the HIGH-condition evaluation. These cut-offs were predefined practical cut-offs for confidence-based grouping and were not selected by data-driven threshold optimization. We first inspected the model-wise distributions of self-reported positive probabilities under prompt 2 (Supplementary Figure 1(b); Supplementary Table 1). The distributions showed that the use of self-reported probability values differed across models. Many models with high classification performance tended to output extreme values such as p = 0.05 or p = 0.95, whereas models with lower classification performance relatively more often produced intermediate or boundary values such as p = 0.40, p = 0.60, or p = 0.75. This pattern suggested that excluding intermediate and boundary probability values could be a reasonable practical approach for identifying predictions with stronger model-reported confidence in this task.

To test this confidence-based grouping, we computed the F1 scores under two conditions: ALL (all projects) and HIGH (only projects with high-confidence). The results obtained using prompt 2 are shown in Figure 4 and Table 3. Under the ALL condition, the precision, recall, and F1 scores were computed for all 150 projects. Under the HIGH condition, samples with 0.25 ≤ p ≤ 0.75 were treated as low-confidence or intermediate predictions and excluded, and the same metrics were recomputed using only samples with p < 0.25 or p > 0.75.

**Figure 4:**
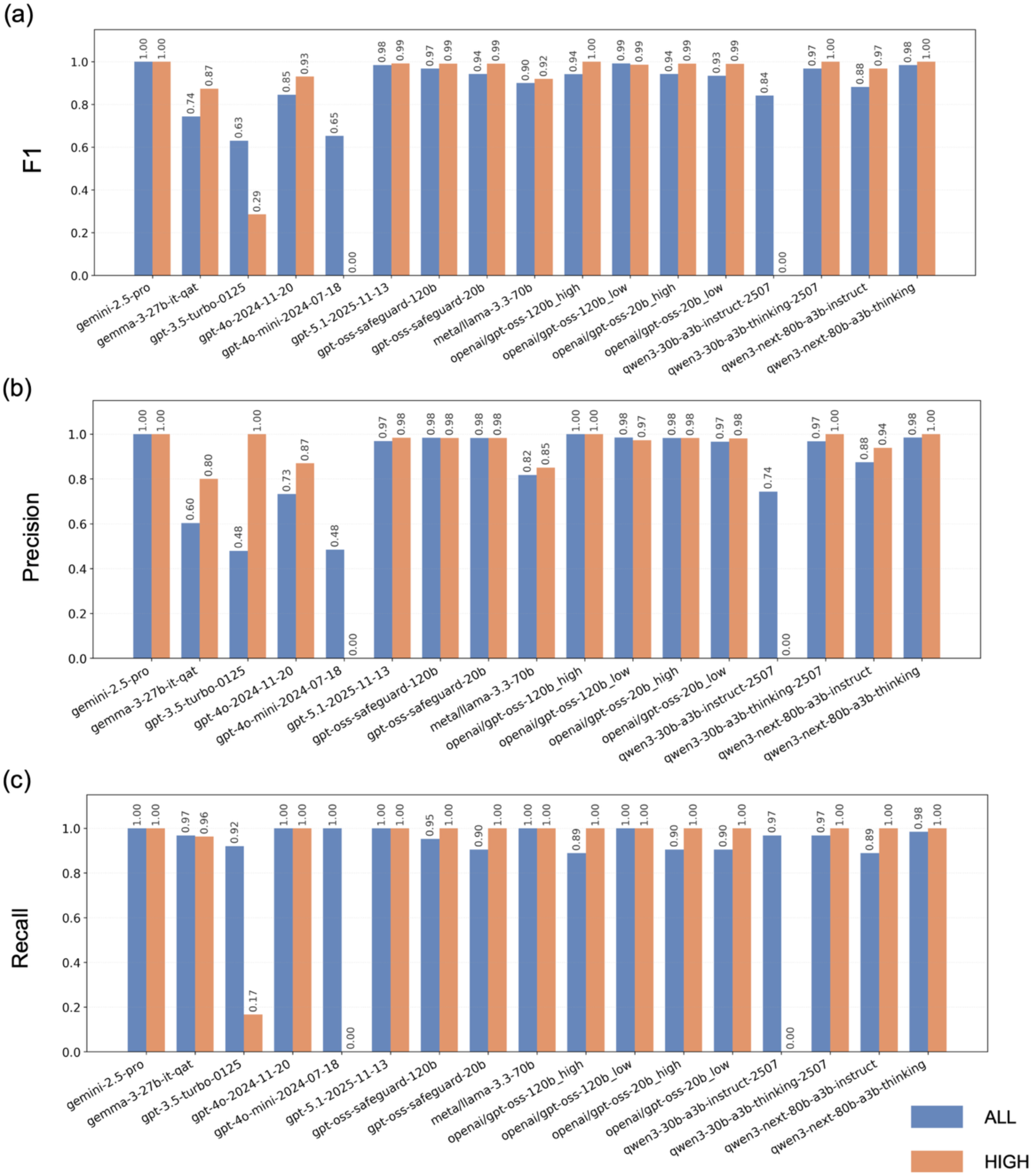
Effects of confidence-based thresholding on classification performance. (a) Comparison of F1 scores under the ALL condition and the HIGH condition (excluding the range 0.25 ≤ p ≤ 0.75) for prompt 2. (b) Comparison of precision under the All and HIGH conditions for prompt 2. (c) Comparison of recall under the All and HIGH conditions for prompt 2.

**Table 3:** Performance comparison between the ALL and HIGH conditions using self-reported probabilities. Precision, recall, and F1 score recomputed after excluding ambiguous predictions (0.25 ≤ p ≤ 0.75). To quantify the trade-off between coverage and reliability across models, the remaining sample size (n) after exclusion is reported.

| Model | Condition | n (ALL) | n' (HIGH) | Precision | Recall | F1 |
| --- | --- | --- | --- | --- | --- | --- |
| gemini-2.5-pro | ALL | 150 | - | 1.000 | 1.000 | 1.000 |
| gemini-2.5-pro | HIGH | - | 143 | 1.000 | 1.000 | 1.000 |
| gemma-3-27b-it-qat | ALL | 150 | - | 0.604 | 0.968 | 0.744 |
| gemma-3-27b-it-qat | HIGH | - | 111 | 0.800 | 0.963 | 0.874 |
| gpt-3.5-turbo-0125 | ALL | 150 | - | 0.479 | 0.921 | 0.630 |
| gpt-3.5-turbo-0125 | HIGH | - | 30 | 1.000 | 0.167 | 0.286 |
| gpt-4o-2024-11-20 | ALL | 150 | - | 0.733 | 1.000 | 0.846 |
| gpt-4o-2024-11-20 | HIGH | - | 116 | 0.870 | 1.000 | 0.930 |
| gpt-4o-mini-2024-07-18 | ALL | 150 | - | 0.485 | 1.000 | 0.653 |
| gpt-4o-mini-2024-07-18 | HIGH | - | 20 | 0.000 | 0.000 | 0.000 |
| gpt-5.1-2025-11-13 | ALL | 150 | - | 0.969 | 1.000 | 0.984 |
| gpt-5.1-2025-11-13 | HIGH | - | 114 | 0.984 | 1.000 | 0.992 |
| gpt-oss-safeguard-120b | ALL | 150 | - | 0.984 | 0.952 | 0.968 |
| gpt-oss-safeguard-120b | HIGH | - | 135 | 0.983 | 1.000 | 0.991 |
| gpt-oss-safeguard-20b | ALL | 150 | - | 0.983 | 0.905 | 0.942 |
| gpt-oss-safeguard-20b | HIGH | - | 140 | 0.983 | 1.000 | 0.991 |
| meta/llama-3.3-70b | ALL | 150 | - | 0.818 | 1.000 | 0.900 |
| meta/llama-3.3-70b | HIGH | - | 116 | 0.851 | 1.000 | 0.919 |
| openai/gpt-oss-120b_high | ALL | 150 | - | 1.000 | 0.889 | 0.941 |
| openai/gpt-oss-120b_high | HIGH | - | 134 | 1.000 | 1.000 | 1.000 |
| openai/gpt-oss-120b_low | ALL | 150 | - | 0.984 | 1.000 | 0.992 |
| openai/gpt-oss-120b_low | HIGH | - | 116 | 0.973 | 1.000 | 0.986 |
| openai/gpt-oss-20b_high | ALL | 150 | - | 0.983 | 0.905 | 0.942 |
| openai/gpt-oss-20b_high | HIGH | - | 137 | 0.983 | 1.000 | 0.991 |
| openai/gpt-oss-20b_low | ALL | 150 | - | 0.966 | 0.905 | 0.934 |
| openai/gpt-oss-20b_low | HIGH | - | 132 | 0.981 | 1.000 | 0.990 |
| **qwen3-30b-a3b-instruct-2507** | ALL | 150 | - | 0.744 | 0.968 | 0.841 |
| **qwen3-30b-a3b-instruct-2507** | HIGH | - | 35 | 0.000 | 0.000 | 0.000 |
| **qwen3-30b-a3b-thinking-2507** | ALL | 150 | - | 0.968 | 0.968 | 0.968 |
| **qwen3-30b-a3b-thinking-2507** | HIGH | - | 88 | 1.000 | 1.000 | 1.000 |
| **qwen3-next-80b-a3b-instruct** | ALL | 150 | - | 0.875 | 0.889 | 0.882 |
| **qwen3-next-80b-a3b-instruct** | HIGH | - | 60 | 0.939 | 1.000 | 0.968 |
| **qwen3-next-80b-a3b-thinking** | ALL | 150 | - | 0.984 | 0.984 | 0.984 |
| **qwen3-next-80b-a3b-thinking** | HIGH | - | 137 | 1.000 | 1.000 | 1.000 |

The number of projects remaining under the HIGH condition, denoted by n′, varied substantially across models, and models that frequently used intermediate probabilities yielded fewer projects n′. Among the models that showed high classification accuracy under the ALL condition, openai/gpt-oss-120b_high, qwen3-30b-a3b-thinking-2507, and qwen3-next-80b-a3b-thinking achieved precision, recall, and F1 scores of 1 under the HIGH condition, with 134, 88, and 137 projects remaining for evaluation, respectively. This implies that the high-confidence decisions self-reported by these models perfectly match the ground-truth labels, suggesting that self-reported probabilities may function as indicators of the reliability of LLM outputs.

In contrast, gpt-3.5-turbo-0125 and gpt-4o-mini-2024-07-18 showed relatively low F1 scores under the ALL condition (0.6304 and 0.6528, respectively); under the HIGH condition, the number of evaluated projects decreased substantially to 30 and 20, respectively. Moreover, even when restricted to the HIGH condition, gpt-3.5-turbo-0125 showed an F1 score of 0.2857 (precision, 1.0000; recall, 0.1667), suggesting that, even in cases where the model self-reported strong confidence, consistency with the ground truth may not be sufficient. For gpt-4o-mini-2024-07-18, the precision, recall, and F1 scores were all zero under the HIGH condition, indicating that in this model, the self-reported probabilities did not adequately reflect the reliability of its decisions.

Thus, rather than showing that self-reported probabilities are always effective, these results can be interpreted as showing that, for models with sufficiently high performance, self-reported probabilities may practically function as reliability indicators. In other words, for high-performance models, an operational design such as “prioritize highly confident decisions, and route ambiguous cases near p = 0.5 for human review” may be feasible.

### AUPRC analysis based on self-reported positive probabilities

In addition to the binary labels, we constructed precision–recall curves using the values that each model self-reported as the probability that each project was positive, and calculated the AUPRC for each model (Supplementary Figure 1(c), Supplementary Table 2). The AUPRC values ranged from 1.000 for gemini-2.5-pro to 0.490 for gpt-3.5-turbo-0125. We further evaluated the correspondence between AUPRC and the F1 score calculated from the binary outputs for all 150 projects under prompt 2. The model rankings based on AUPRC showed strong agreement with the rankings based on the all-sample F1 scores. Specifically, in 12 of the 17 models, the difference between the F1 rank and the AUPRC rank was within ±1, and four of the top five models were shared between the two metrics (Supplementary Table 2). To visualize this relationship, we added a horizontal bar plot comparing the F1 score and AUPRC for each model as Supplementary Figure 1(d), allowing readers to inspect the correspondence between the magnitude and ranking of the two metrics. This agreement indicates that the model-level conclusions derived from the all-sample F1 scores are broadly supported by the AUPRC evaluation based on self-reported positive probabilities. However, the self-reported positive probabilities were not distributed as precise continuous probability values. Instead, they were concentrated mainly at seven discrete values: 0.05, 0.40, 0.60, 0.75, 0.80, 0.90, and 0.95 (Supplementary Table 1). This pattern may reflect the probability guidance provided in the prompt. The scripts used to generate the AUPRC and confidence-distribution analyses is provided in Supplementary File 1.

### Reproducibility of LLM outputs under repeated inference

Across the five independent repeated runs, the binary labels and self-reported positive probabilities were identical for every evaluated project within each model. The per-accession results and reproducibility summaries are provided in Supplementary Table 3 and the reproducibility supplement is provided in Supplementary File 2. This result indicates that, under the tested settings and execution session, both openai/gpt-oss-120b_low and qwen3-next-80b-a3b-thinking produced stable outputs for the n = 50 reproducibility subset. As a complementary check, we compared the original single-run outputs used for Table 1 with the outputs from the 5-run reproducibility experiment on the same n = 50 subset (Supplementary Table 4). The binary labels matched for all 50 projects in both models. The self-reported probabilities also matched for all 50 projects in qwen3-next-80b-a3b-thinking. In openai/gpt-oss-120b_low, however, the self-reported probability differed for 5 of 50 projects: four cases changed from 0.95 in the original run to 0.75 in the reproducibility run, and one case changed from 0.75 to 0.80. These shifts occurred within the high-confidence positive probability range defined by the prompt and did not cross the 0.5 decision boundary; therefore, they did not affect the binary classification metrics reported in Table 1.

### Comparison of the time required to process LLM outputs

In addition to classification accuracy, processing speed is also important when handling large volumes of data. We therefore compared the per-task execution time of each model and summarized the runtime measurements in Supplementary Figure 2 and Supplementary Table 5.

Because closed models are executed on high-performance servers, their processing time tends to be shorter than those of models executed locally. In contrast, when comparing speeds among open-weight models released in 2025, three factors must be considered: (1) whether the model is a reasoning model (i.e., whether it is designed to use the thinking mode), (2) the reasoning effort setting in gpt-oss models, and (3) differences in architecture design, such as dense versus mixture of experts (MoE). Among the open-weight models executed locally, smaller-parameter models tended to have shorter execution times. Additionally, settings that do not include a reasoning (reasoning/thinking) process or allocate less time to reasoning tend to produce outputs more quickly. Conversely, when a reasoning process was included or more time was allocated to reasoning, the execution time increased compared to models of the same scale that did not perform reasoning. For example, when comparing the execution time between Qwen3 thinking models and instruct models, the thinking models showed higher performance, but required substantially longer execution times. Focusing on gpt-oss 20b and gpt-oss 120b in both models, the execution time was longer when the reasoning effort was high than when it was low. Moreover, comparing gpt-oss 20b with high reasoning effort to gpt-oss 120b with low reasoning effort, the latter completed processing in a shorter time despite its larger model size, because the reasoning effort was lower. Furthermore, MoE-type open-weight models, which were frequently observed among the 2025 releases (gpt-oss series, Qwen3 series), showed improved execution efficiency compared with conventional dense open-weight models, such as Llama 3.3 and Gemma 3 27 B. For instance, gpt-oss-120b/20b were designed as MoE-based open-weight models with a total parameter count of approximately 117 B/21 B. However, during inference, only a subset of the parameters selected according to the input becomes active, enabling faster execution relative to the overall model size ^17^. Similarly, Qwen3 adopts a highly sparse MoE design wherein, despite having 30 B or 80 B parameters, only approximately 3 B parameters are selectively used; moreover, an improved processing speed relative to older Qwen models has been reported ^18^.

The results of this experiment showed that, while Llama 3.3 and Gemma 3 27 B did not include a reasoning process, MoE-type Qwen instruction models without reasoning and gpt-oss with reduced reasoning (Low) completed processing substantially faster than conventional dense models.

Moreover, even for Qwen thinking models that perform reasoning and gpt-oss with increased reasoning (High), higher output accuracy was achieved while maintaining the execution efficiency advantage of the MoE designs, with execution times comparable to or shorter than those of conventional models. This indicates that open-weight models have made notable progress in terms of accuracy and processing speed over a short period.

### F1–runtime trade-off in local open-weight model execution

To evaluate the practical balance between classification accuracy and processing speed, we compared the F1 score and mean per-project runtime for ten locally executed open-weight conditions. The source data are provided in Supplementary Table 6, and the corresponding scatter plot is shown in Supplementary Figure 2(e). In the Qwen3 family, switching from the instruct to the thinking variant improved F1 from 0.841 to 0.968 for Qwen3 30B and from 0.882 to 0.984 for Qwen3 Next 80B, but increased mean runtime from 4.2 to 24.9 s/project and from 5.0 to 38.3 s/project, respectively. In the gpt-oss family, increasing reasoning effort from low to high increased runtime by approximately 4.8-fold for the 20B model and 3.8-fold for the 120B model, but did not improve the F1 score to the same extent. The dense reference models were less favorable in this trade-off space: meta/llama-3.3-70b required 87.2 s/project with F1 = 0.900, whereas gemma-3-27b-it-qat achieved F1 = 0.744 at 20.8 s/project.

### Extraction and summarization of project and sample information from metadata

In addition to the classification task above, the workflow created in this study implements a function that allows users to specify arbitrary column headers for the retrieved sample metadata and organize the information as tabular data. Using the metadata of projects judged as positive by the LLM in the classification step as input, the workflow automatically generates a table, as shown in Supplementary Table 7.

For example, when items such as sample genotype, tissue name, treatment method, treatment concentration, and treatment duration were specified, the corresponding information was extracted from the descriptions in the metadata and output. Contents not described are output as “Unspecified.” Thus, the model can be flexibly instructed on what to extract by simply listing the desired items in the prompt. Previously, rule-based methods relied on dictionaries prepared on the program side or fixed keyword matching. These methods could not sufficiently handle variations in descriptions and the diversity of expressions, and were not flexible with respect to adding new items or changing definitions. As a result, flexible and simple information extraction, such as that demonstrated here, was not easy in terms of both coverage and extraction accuracy; however, the use of LLMs significantly mitigated this issue. However, the accuracy of this detailed information output was not verified or evaluated in this study. Unlike the binary classification evaluated above, the output format is diverse and complex, making the evaluation difficult. Although LLMs make a variety of outputs possible, the creation of ground-truth datasets and the speed of human evaluation have not kept pace with the progress in LLM development. Although generating such complex natural language outputs has become technically feasible owing to improvements in LLM capabilities, developing methods to evaluate these outputs remains a challenge.

## DISCUSSION

In this study, we applied LLMs to an initial metadata screening task that has traditionally been handled by humans or has not been sufficiently addressed because of large target volumes. We highlighted three major points of contention when using LLMs to substitute such work: (1) accuracy, (2) adoption and operational cost (both financial cost and human effort), and (3) processing speed. Using a classification task on public RNA-seq metadata, we compared multiple LLMs under identical conditions and examined the practicality of LLM-based curation support from the perspectives of these three axes.

In this study, we defined a baseline (“Keyword Search Only”) in which all projects returned by keyword search are treated as positive and compared LLM classification performance against it. Because the baseline retrieved all hits, the recall was 1.00; however, precision was 0.42, and the F1 score was 0.59, indicating that many false positives were included. This means that in dataset discovery for meta-analysis and reanalysis, simple keyword search alone allows a large number of projects to be mixed in that contain the search terms but do not satisfy the target conditions (e.g., ABA-treated RNA-seq samples, target organism, presence of controls). Tools such as GEOmetadb and pysRADB are useful for accessing, querying, and organizing public GEO/SRA metadata, and they correspond to the retrieval and metadata collection stages of workflows such as ours. However, these tools do not by themselves directly replace the downstream semantic eligibility judgment evaluated in this study. The main bottleneck addressed here was therefore not simply retrieving records that contain relevant query terms, but determining whether the retrieved project-level and sample-level metadata satisfy user-defined biological criteria, such as the presence of exogenous ABA-treated *Arabidopsis* RNA-seq samples together with corresponding untreated controls. LLM-based classification substantially improved the accuracy, precision, and F1 score of many models.

Among the closed models, gemini-2.5-pro (prompt 2) exhibited the best performance, and gpt-5.1 (prompt 2) achieved a high F1 score of 0.98 (Table 1). The open-weight models also demonstrated high performance; for example, gpt-oss-120b_low (prompt 2) achieved an F1 score of 0.99, and qwen3-next-80b-a3b-thinking (prompt 2) achieved an F1 score of approximately 0.98. These results suggest that even without using the latest closed models, the currently released mid-scale open-weight models can potentially execute screening support tasks such as criterion-based life science metadata classification with high accuracy. Another notable point is that, as of 2025, multiple open-weight models outperform the gpt-4o series widely used in 2024. This suggests that open-weight models are improving rapidly. When discussing accuracy, it is necessary to clarify that the magnitude of the F1 score as well as the FPs and FNs constitute a large practical burden. If FPs occur frequently, the workload increases during the downstream curation steps. If FNs are frequent, useful projects are increasingly missed, potentially reducing the coverage of the datasets required for comprehensive exploration and reanalysis. Therefore, by creating and checking both a prompt designed for recall (to reduce missed positives) with concise criteria, and a prompt designed for precision (to reduce FPs) with detailed criteria, data discovery can be conducted while suppressing both missed hits and unnecessary candidate contamination. Because accuracy is shaped not only by model choice but also by instructions, we then examined how prompt design shifts the precision/recall trade-off across models. In this study, we compared two prompts: prompt 1 tried to avoid missing relevant projects, and prompt 2 tried to reduce false positives. Indeed, prompt 2 tended to improve precision in many models and contributed to the suppression of FPs (Table 2). However, this effect was not uniform across models. In some models, prompt 2 was accompanied by a decrease in recall. This suggests that the prompt can push the decision threshold in the direction of avoiding misdetections, leading to more cautious judgments and reducing positive calls.

Conversely, there were cases in which prompt 2 improved recall while maintaining precision. These findings indicate that, even with prompts designed with the same intent, the effect differs depending on the internal representations and training characteristics of the model. Thus, a prompt design can influence LLM outputs. However, the degree of control achieved by prompts varies according to the model type. Accordingly, after clarifying whether FPs or FNs should be prioritized for the target task, it is necessary to test and evaluate the prompt in the intended model to confirm that it functions as designed. This analysis should not be interpreted as a comprehensive prompt-engineering study.

Instead, it was designed to test whether changing the stringency of task-specific eligibility criteria, including conservative handling of insufficient evidence, could shift the precision-recall balance. Because the effects of prompt design depend on the target task, metadata characteristics, and the desired balance between false positives and false negatives, it is unlikely that a single universally optimal prompt will be suitable for all curation settings. Therefore, in practical applications, prompts should be designed and empirically tested according to the specific objective, data type, and expected sources of ambiguity in each use case. Based on the results of this verification, we offer the following recommendations to researchers and practitioners designing similar metadata-screening tasks. First, define explicit positive and negative criteria for the target task, decide a priori whether false positives or false negatives are more problematic, and design the prompt accordingly; in practice it can be useful to compare a recall-oriented prompt with concise inclusion criteria and a precision-oriented prompt with more detailed exclusion criteria, as we did with prompt 1 and prompt 2 in this study. Second, with downstream automated parsing in mind, explicitly specify a structured output format such as JSON and, where appropriate, request a self-reported confidence or probability score alongside the binary label. Third, before applying the workflow at scale, validate both the prompt and the model on a small manually labeled subset; because the optimal prompt depends on the data and target, comparing several candidate prompts on a small dataset and selecting based on observed accuracy is more practical than fine-tuning the wording in isolation. Finally, given the rapid progress of recent LLMs, we expect that, in order to achieve the intended objective, clearly specifying the task objective, decision criteria, and expected output format will become increasingly important, rather than seeking further improvements in accuracy through fine-grained prompt-engineering techniques.

Beyond accuracy, practical deployment also depends on resource costs, so we compared manual-only curation with closed-model APIs and locally executed open-weight models. Here, we define the term “cost” broadly as the resources required for researchers to accomplish curation tasks (such as fees, time, and effort). In manual-only workflows, researchers must read metadata (and sometimes related papers) and judge eligibility project by project, which becomes impractical at scale due to accumulated time, labor costs, and cognitive burden. Although adopting LLMs introduces additional operational costs (e.g., API fees or local setup and pipeline development), these can be weighed against the reductions and benefits enabled by LLM-based workflows. Considering the above costs, LLM adoption can significantly reduce many costs humans have traditionally borne. In the workflow constructed in this study, by automating metadata retrieval, preprocessing, LLM-based classification, and results aggregation, researchers no longer need to carefully read all projects individually; in similar tasks in the future, researchers can focus on final confirmation after the LLM narrows down the candidates. This is not merely doing the same work faster; rather, it is a qualitative shift that makes exploration and organization tasks feasible within realistic time and effort—even at scales that were previously too large to curate. Additionally, when operating open-weight models locally, a specific model version that has been benchmarked for accuracy can be fixed and used under the researcher’s control while maintaining a low long-term financial cost. Although closed-model APIs may change output tendencies or performance without notice owing to provider updates, local open-weight models enable continued use of the same version, which is advantageous for research reproducibility; similar output tendencies and performance levels can be more readily maintained across reanalysis and replication performed at different times. However, fixing the model version alone does not guarantee complete output reproducibility. Even under identical prompt and temperature settings, outputs may be affected by inference-time randomness, software implementation, inference backend, hardware-dependent numerical differences, and session-specific execution conditions. Therefore, local open-weight models should be interpreted as improving version stability and supporting reproducible evaluation, rather than guaranteeing complete determinism of all generated outputs. Therefore, “manual-only,” “closed-model use,” and “local open-weight use” each have different cost structures and benefits.

We also evaluated processing speed, focusing on how model architecture and inference settings influence execution time in large-scale curation. Processing speed strongly affects the practicality of large-scale metadata curation. Compared with manual visual inspection, batch processing by LLMs scales more effectively as the number of projects increases and can substantially shorten the stages of discovery and condition-based filtering. Our measurements showed clear differences in processing time across models (Supplementary Figure 2, Supplementary Table 5). Closed models tend to exhibit shorter mean processing times because they operate on high-performance servers. In contrast, for locally executed models, the speed was strongly influenced by differences in the model architecture and inference modes. For example, Qwen3 thinking models (designed to perform reasoning) showed high classification performance, but tended to require longer execution times than instruction models. Similarly, even within the same gpt-oss model, setting the reasoning effort to high substantially increased the runtime compared to that with low reasoning effort, indicating that the level of reasoning is a major determinant of speed. Furthermore, many open-weight models released in 2025 adopt MoE designs, which tend to improve execution efficiency compared with dense models; this trend was also confirmed in our results. Hence, speed depends strongly not only on model size but also on the actual computational amount used during inference. Therefore, when selecting models, it is important to consider speed requirements in addition to accuracy, in accordance with the number of items to process and the operational constraints. The F1–runtime comparison among locally executed open-weight models further supports this point, suggesting that practical model selection for large-scale metadata screening should consider both classification performance and local throughput (Supplementary Table 6 and Supplementary Figure 2(e)). In particular, low-reasoning gpt-oss settings provided favorable F1–runtime trade-offs, whereas thinking or high-reasoning settings increased runtime substantially and did not always yield proportional gains in F1. These findings suggest that, depending on the scale of the screening task and the acceptable balance between false positives and false negatives, faster lower-reasoning settings may be suitable for initial candidate filtering, while slower reasoning-oriented settings may be useful when higher-confidence judgments are required or when the number of candidate projects is limited. At the same time, the observed trade-off also highlights the recent progress of MoE-based open-weight models. Compared with dense reference models, recent MoE-based models achieved stronger combinations of F1 score and runtime under the evaluated local execution conditions. This suggests that architectural improvements, together with adjustable reasoning-effort settings, may make locally executable open-weight models increasingly practical for metadata-screening workflows that require both reliable classification and scalable processing speed.

Taken together, these findings motivate an operational design that explicitly balances accuracy, cost, and speed rather than always using a single highest-accuracy model. Based on these results, when introducing LLMs to handle a number of targets that are difficult for humans to confirm manually, it may be more effective to design trade-offs among accuracy, speed, and cost rather than always using a single highest-accuracy model. For example, a staged design is possible that prioritizes recall (avoid missing positives) during initial discovery, prioritizes precision (reduce FPs) during subsequent screening, and finally confirms the resulting outputs. In this study, we also attempted to have the models output a positive probability alongside binary classification, which could potentially be used to extract borderline cases that should be routed for human review or to select candidates to pass into a second-stage process. Indeed, among the models that showed high classification accuracy under ALL conditions, gpt-oss-120b_high, qwen3-30b-a3b-thinking-2507, and qwen3-next-80b-a3b-thinking achieved a precision, recall, and F1 of 1.00 under the HIGH condition, covering 134, 88, and 137 projects, respectively. This suggests that self-reported probabilities can serve as a useful proxy for reliability, at least for classification tasks. Accordingly, an operational design may become realistic in which samples with intermediate probabilities are routed to humans, whereas those with high confidence probabilities are processed automatically.

However, this trend does not apply to all models. For example, gpt-3.5-turbo-0125 and gpt-4o-mini-2024-07-18 showed relatively low F1 scores under the ALL condition (0.6304 and 0.6528, respectively). Under the HIGH condition, the number of evaluated projects dropped to 30 and 20, respectively, with substantially reduced metrics in some cases. Therefore, it is important to note that a high confidence does not necessarily guarantee accuracy. When incorporating model confidence into operations, it is desirable to first confirm whether similar behavior is observed for the chosen model and task and to use confidence in combination with human review.

In this study, to complement the F1 scores calculated from binary outputs across all samples, we constructed precision–recall curves from each model’s self-reported positive probabilities and calculated AUPRC. The model rankings based on AUPRC showed strong agreement with the rankings based on the F1 scores calculated from all 150 projects under prompt 2. This indicates that the main conclusions of our model comparison based on F1 scores are also supported by an evaluation using self-reported positive probabilities. At the same time, the self-reported positive probabilities did not behave as precise continuous probability values. Across the 17 models, they were concentrated at a small number of discrete values, mainly 0.05, 0.40, 0.60, 0.75, 0.80, 0.90, and 0.95 (Supplementary Table 1). This likely reflects the probability guidance provided in the prompt. Therefore, the AUPRC values reported in this study should not be interpreted as metrics based on precise continuous probability values. Rather, they should be interpreted as summary metrics based on the ordering information contained in discrete confidence scores guided by the prompt. Nevertheless, the strong agreement between AUPRC rank and the all-sample F1 rank suggests that self-reported positive probabilities may be useful, at least for some high-performing models, for ranking candidate projects in metadata screening or for prioritizing high-confidence cases for automated processing. However, because self-reported positive probabilities do not necessarily reflect reliability equally well across all models, their relationship with the actual ground-truth labels should be validated in advance for each target model and task before they are used operationally.

The confidence-based analysis should be interpreted with caution because the 0.25 and 0.75 cut-offs were predefined practical cut-offs for confidence-based grouping, not thresholds optimized from the data. The model-wise distributions of self-reported probabilities showed that many models with high classification performance tended to output extreme values such as p = 0.05 or p = 0.95, whereas models with lower classification performance relatively more often produced intermediate or boundary values such as p = 0.40, p = 0.60, or p = 0.75 (Supplementary Figure 1(b), Supplementary Table 1). These observations suggest that, although not optimized, the predefined 0.25 / 0.75 cut-offs provided a reasonable operational grouping for this task.

Although the repeated inference experiment showed stable outputs within the tested execution session, this result does not guarantee complete determinism across all models, hardware environments, inference backends, or execution sessions. Consistent with this caveat, the comparison between the original Table 1 run and the later reproducibility experiment showed a small cross-session drift in self-reported probabilities for openai/gpt-oss-120b_low, while qwen3-next-80b-a3b-thinking showed no such difference. The observed shifts occurred within the high-confidence positive probability range defined by the prompt and did not cross the 0.5 decision boundary, so the binary classification metrics were unaffected; however, the number of projects included in the HIGH-confidence subset could change by a few projects if recomputed in a different session. These observations indicate that the self-reported probabilities should be interpreted as useful confidence signals rather than exact probability values that are guaranteed to be identical across sessions.

Finally, we note the main limitations of the present evaluation and outline directions for future work. The first limitation of this study is that the evaluated task was a binary classification, and we did not sufficiently evaluate the accuracy of more complex structured outputs (e.g., extracting genotype, tissue, treatment concentration, and treatment duration). While LLMs can produce diverse output formats, the more complex the output, the more difficult it is to create ground-truth annotations and design evaluation procedures, making it difficult for evaluation methods to keep pace with rapid advances in output generation. Developing evaluation methods for complex information extraction tasks is an important direction for future research.

The second limitation is that the LLM decisions are constrained by the information contained in the public metadata provided as input. This benchmark evaluates semantic classification under fixed input conditions. Information that was not retrieved by the keyword search or not included in the integrated metadata text was outside the scope of the present evaluation. Tasks performed by human curators—cross-checking multiple sources such as project pages, sample descriptions; the full text of associated papers, and supplementary materials; and resolving inconsistencies or filling in missing information—are not included in this workflow. Consequently, when the metadata is incomplete, ambiguous, or inconsistent with the actual experimental content, an LLM may produce a positive or negative decision that is consistent with the provided metadata, even if the true dataset content differs.

Therefore, although this workflow is effective as an initial screening step to narrow candidates, final inclusion decisions and borderline cases should be verified by humans using additional sources (e.g., related pages and papers). The same limitation also applies to self-reported confidence scores: when the input metadata are incomplete or ambiguous, high confidence does not necessarily guarantee that the decision reflects the true experimental content. To mitigate this limitation, it is important to design how to collect, integrate, and present the information necessary for LLM decisions, such as which sources to use and at what scope and granularity to construct an appropriate input context.

Third, the dataset evaluated in this study was derived from search results based on a specific organism, treatment condition, and data type (*Arabidopsis thaliana*; ABA treatment; bulk RNA-seq), and the number of projects available for evaluation was limited (n = 150). In this pilot evaluation, bulk RNA-seq was intentionally selected as a tractable initial test case because project-level eligibility criteria could be clearly defined and a reliable ground-truth dataset could be constructed. In contrast, single-cell RNA-seq, paired bulk and single-cell RNA-seq datasets, and multi-omics datasets are more complex because they require additional information to interpret the samples correctly, such as cell type, experimental method, batch information, and sample relationships across datasets or modalities. Additional validation is required to determine whether similar performance and model rankings are maintained under different organisms, treatments, metadata qualities, and retrieval strategies. In particular, the present evaluation was performed only on the candidate pool retrieved using the keyword search strategy defined in this study. If different keywords or search strategies are used, the retrieved candidate projects may change, and this could affect the downstream LLM performance evaluation and model ranking. Evaluating the workflow under alternative retrieval strategies is therefore an important direction for future work.

The workflow is designed to be configurable, but the performance estimates reported here should be interpreted as specific to the evaluated *Arabidopsis* ABA-treatment bulk RNA-seq screening task.

Applying the workflow to other organisms, treatments, data types, or databases requires task-specific criteria, appropriate input sources, and independent accuracy validation. Additional validation is therefore required to determine whether comparable performance can be achieved when the workflow is generalized to other organisms, treatment conditions, data types, or domains with substantially different metadata quality and reporting conventions. Accordingly, applications to more complex scenarios would require the incorporation of additional data sources, substantial modifications to the workflow, and purpose-specific accuracy validation.

Fourth, because the benchmark labels were assigned by a single curator, independent multi-curator annotation and inter-annotator agreement were not evaluated.

Nevertheless, we conducted a cross-model comparison using two prompts with different levels of strictness and a diverse set of models from multiple organizations that differed in terms of release timing and performance. We observed both models whose classification performance was largely unaffected by prompt changes and whose performance improved when the prompts were made stricter. In addition, newer models, which are typically considered to perform better, tend to achieve better classification results. Taken together, these results suggest that the results of this study are not specific to a particular model or a single prompt setting but rather have a certain generality within this task setting.

Overall, this study shows that LLM-based classification can substantially improve keyword-search baselines in the practical task of exploring public metadata and that high accuracy is becoming achievable even with open-weight models. As model performance continues to improve and usage barriers decline, what becomes critical in research is not only the model’s raw capability, but also the end-to-end design on the user side: for what purpose, what input to provide, what output to obtain, and how to validate and operate the system. The workflow presented here provides a concrete framework for design and validation and serves as a foundation for further extensions toward automated metadata screening and early-stage curation.

## METHODS

In this study, we used LLMs to perform project-level binary classification on publicly available RNA-seq projects to determine whether each project contained both (i) exogenous abscisic acid (ABA)-treated samples in *Arabidopsis thaliana* and (ii) the corresponding untreated control samples. The workflow consisted of the following three steps. First, we retrieved the metadata from public databases and processed them for downstream analysis. Second, we used an LLM to classify the retrieved metadata according to predefined criteria. Third, we validated the LLM outputs by comparing them with a human-labeled ground-truth dataset. The workflow scripts developed and used in the study are publicly available on GitHub (https://github.com/mshintani22/open-weight-llm-metadata-curation-workflow).

### Retrieval of public RNA-seq metadata using APIs

To retrieve metadata from public databases, we queried BioProject and GEO via NCBI Entrez Programming Utilities (E-utilities) using keyword-based search queries and obtained the accession IDs for projects that matched the search conditions. For the GSE accessions obtained from GEO, we first used ELink (E-utilities) to map them to the corresponding BioProject records. When direct mapping could not be obtained, we attempted two alternative routes: (i) starting from GEO DataSets records and tracing to BioProject via SRA records, and (ii) tracing to BioProject via BioSample records. If a project could not be mapped through these routes, we supplemented the information using the TogoID API ^19^ and GEO MINiML (XML) and normalized the identifiers to BioProject IDs.

For each BioProject, we retrieved both the project overview and sample/run metadata using the European Nucleotide Archive read_run API. A comprehensive database search was performed using queries containing the following keywords. For GEO: ("Expression profiling by high throughput sequencing"[Filter]) AND ("Arabidopsis thaliana"[porgn]) AND ("ABA"[All Fields] OR "abscisic acid"[All Fields]); for BioProject:("Arabidopsis thaliana"[Organism]) AND ("transcriptome gene expression"[Filter] AND "material transcriptome"[Filter] AND "method sequencing"[Filter]) AND ("ABA"[All Fields] OR "abscisic acid"[All Fields]). The search was executed on December 7, 2025, and yielded 127 GEO and 127 BioProject projects. After converting GEO IDs to BioProject IDs and comparing the results, 104 projects were common to both databases, whereas 23 projects were unique to each database. After the duplicates were removed, 150 projects were used for subsequent analyses. A list of the 150 projects is available in Supplementary Table 8.

The program developed in this study allowed users to replace the organism name (e.g., “*Arabidopsis thaliana*” in the above queries) and the search keywords (e.g., “ABA, abscisic acid”) during runtime. With these inputs, the program automatically constructs search expressions with the same structure as above and performs cross-database searches against GEO and BioProject for any specified organism and treatment/stress condition. Beyond these runtime options, users can add limited task-specific instructions to the prompts and modify the target items used for sample-level extraction. However, the current implementation is coupled to the output formats parsed by the downstream scripts and to the API structures of GEO, BioProject, and ENA; therefore, extension to other databases requires additional database-specific API implementation.

### Retrieved metadata

The project- and sample-level metadata were retrieved separately as (i) project-wide overview information and (ii) information associated with each sample (run). We then integrated these into a single text file so that the LLM could reference both sources within a single input. In the integrated text, the project overview is placed first, followed by a per-sample listing of the sample information indexed by the sample IDs. During integration, information that was redundantly repeated across samples was consolidated into a project-level section, whereas sample-specific information was retained at the sample level. This design reduces the input length while preserving the information required to determine the presence of the treatment and control groups. The resulting integrated metadata text was used as the LLM input in the subsequent steps. The metadata obtained from running the workflow is available in the Supplementary File 4.

### Model types and parameter settings

In this study, we primarily used open-weight models that could be downloaded and executed locally. The open-weight models released by OpenAI, Alibaba, Meta, and Google were used. In addition, to compare the classification performance of the open-weight and closed models, we used the API-based closed models provided by OpenAI and Google. The details of all models are summarized in Table 4. Quantized and converted open-weight models released by OpenAI, Alibaba, Meta, and Google were downloaded from Hugging Face and executed locally using LM Studio, an application on a local PC. For the OpenAI open-weight models that support a reasoning effort setting (gpt-oss-120b and gpt-oss-20b), we tested both High and Low settings. OpenAI and Google’s closed models were run via their APIs from December 14 to 15, 2025. We evaluated ten open-weight models. However, for gpt-oss-120b and gpt-oss-20b, we tested two reasoning effort settings (High and Low), bringing the total number of open-weight evaluation conditions to 12. We then included five closed-source models and used two prompts per condition, resulting in a comparison of outputs across 34 conditions. All models were run with temperature = 0 and maximum tokens = 60,000. Each project was processed as an independent request without carrying over chat history from previous projects. In the initial benchmark, each model was run once per project and prompt condition. The context window size used in this study is sufficient to accommodate the input metadata, prompts, and generated outputs.

**Table 4:**
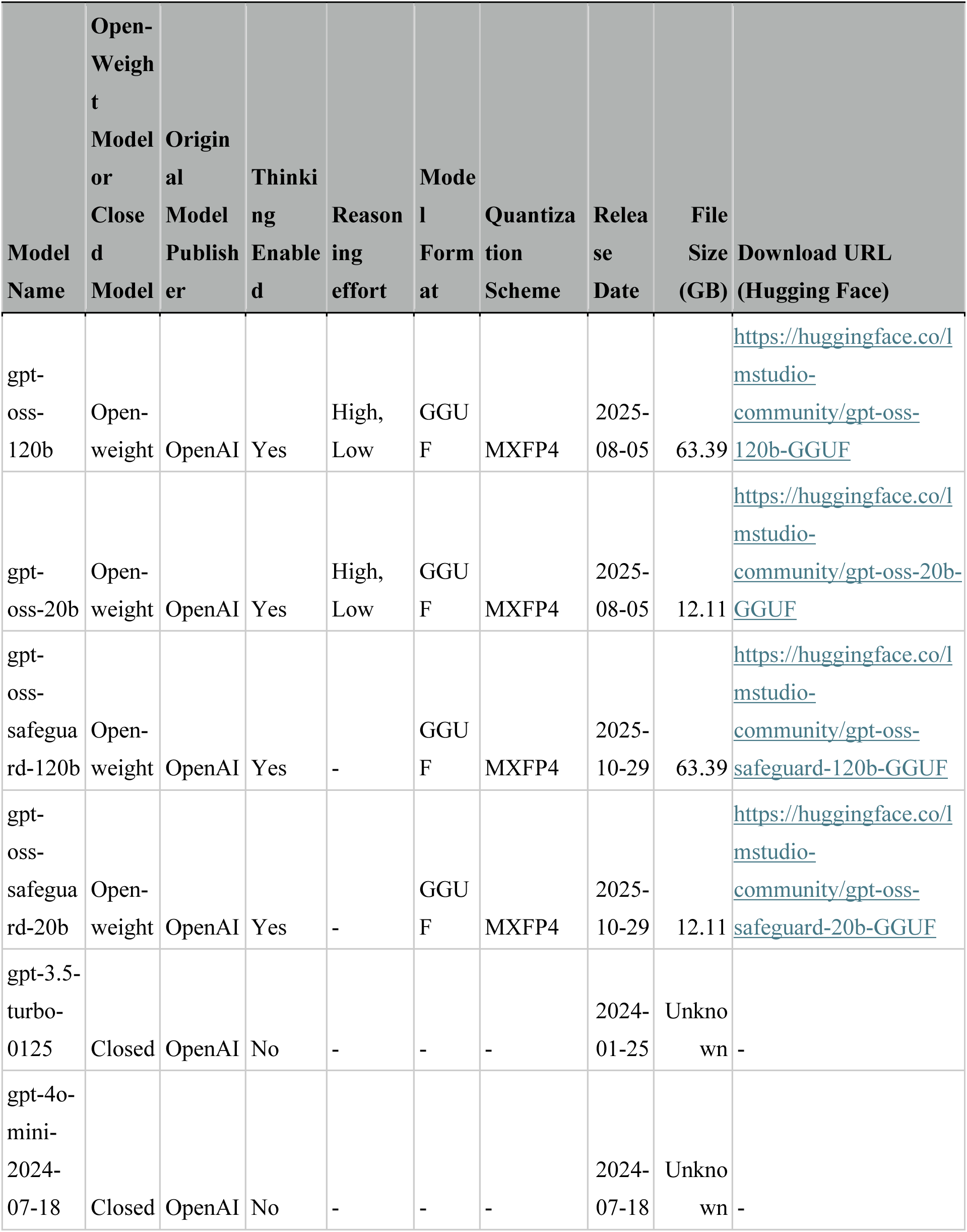
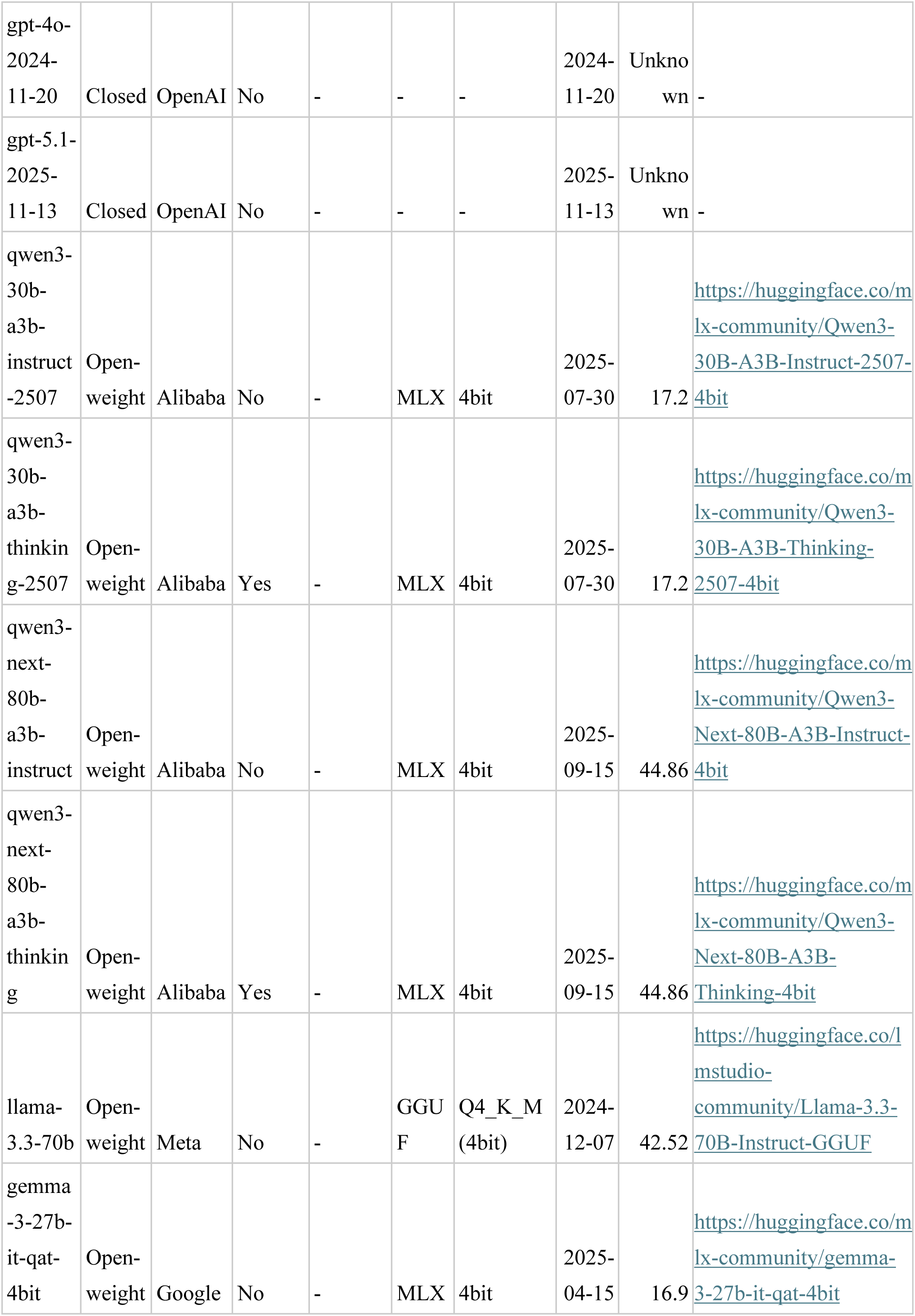
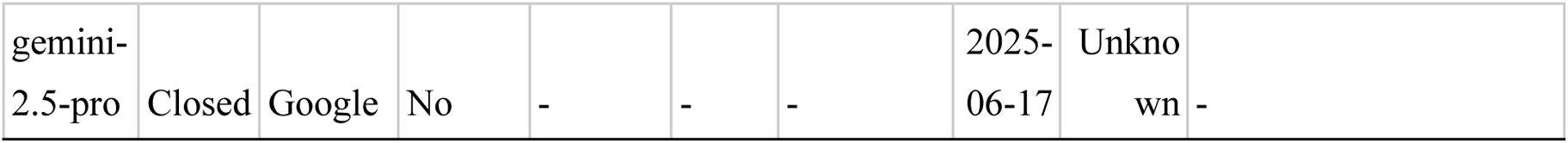
List of language models used in this study. Evaluated models (open-weight, closed), provider, thinking/reasoning settings, model format and quantization, release date, file size, and download sources.

We used the same input metadata and output formats (JSON) for all the models.

### Repeated inference experiment

To assess the reproducibility of model outputs and self-reported probabilities, we performed repeated inference on a subset of 50 projects randomly sampled from the 150-project benchmark using a fixed sampling seed of 42. Two representative open-weight models, openai/gpt-oss-120b_low and qwen3-next-80b-a3b-thinking, were tested under prompt 2 with temperature = 0; the same subset was processed five independent times. No inference-time seed was fixed. We did not repeat the main benchmark under multiple temperature or reasoning settings; instead, the evaluated settings were fixed for each model, and reproducibility was assessed under the same prompt 2 and temperature = 0 condition.

### Tools and hardware used for LLM execution

The open-weight models were executed using LM Studio. These models were downloaded from Hugging Face through LM Studio, loaded into LM Studio, and hosted on a local machine. The program developed in this study repeatedly executes inferences by calling the hosted models using LM Studio’s local HTTP endpoint. We used LM Studio version 0.3.33. The internal tools used were Metal llama.cpp v1.61.0, LM Studio MLX v0.34.0, and Harmony (Mac) v0.3.5. The execution machine used was a Mac Studio (2025 model) with Apple M4 Max, 16-core CPU, 40-core GPU, 128 GB RAM, and 2 TB SSD. The closed models were accessed via OpenAI and Google API keys and executed on provider-managed servers.

### Prompts used for project-level classification and pilot metadata-based sample extraction/summarization

To examine how the strictness of rule definitions in prompts affects the classification performance, we prepared two prompts. Prompt 1 included only the minimum set of positive criteria and rules, prioritizing the reduction of false negatives and emphasizing recall. Prompt 2 added stricter, more detailed conditions to the positive criteria, prioritized the reduction of false positives, and emphasized precision. In particular, prompt 2 instructed the model to require explicit evidence of exogenous ABA treatment and corresponding untreated controls within the same project and to avoid inferring eligibility when the project-level or sample-level metadata were insufficient or ambiguous. In addition, as a pilot feature implemented independently from the binary classification evaluation, we added a function that outputs sample information as a table for projects classified as positive, based on metadata. The prompts used in this study are available in the Supplementary File 5. The outputs from running the metadata search and retrieval scripts, as well as the directories of LLM outputs generated using prompt 1 and prompt 2, are provided in Supplementary File 6.

### Construction of the evaluation dataset and performance assessment

For the 150 projects retrieved by the search, the ground-truth labels were assigned by a single curator according to predefined criteria under the same fixed-input setting used for LLM evaluation. The curator manually inspected only the integrated metadata text retrieved by the workflow and provided to the LLMs, without consulting associated full-text articles, supplementary materials, external web pages, or other external sources. A label of 1 was assigned only when all predefined positive criteria were explicitly satisfied; otherwise, the project was labeled 0. This design allowed us to evaluate semantic classification based on the same metadata input provided to each model, rather than combining classification performance with additional information-retrieval performance. A project was labeled 1 (positive) only when the integrated metadata text explicitly satisfied the following conditions: (i) The project is explicitly described as RNA-seq. (ii) The target species is explicitly *Arabidopsis thaliana*. (iii) Exogenous ABA treatment (“ABA” or “abscisic acid”) is explicitly applied to the RNA-seq samples. (iv) The same project explicitly included untreated control samples corresponding to the ABA-treated samples.

A project was labeled 0 (negative) if any of these conditions were missing. This included cases where “ABA” appeared only in background descriptions without explicit evidence of treatment application, or where ABA-treated samples were present but untreated controls could not be confirmed within the same project, meaning a valid comparative design was not established.

Based on these rules, the evaluation dataset comprises 63 positive (1) and 87 negative (0) results. The ground-truth table (n = 150), including BioProject IDs and manually curated labels, is available in the Supplementary Table 9

We then compared the evaluation labels with the LLM outputs, and defined the confusion matrix components as follows:

- True positive (TP): evaluation label = 1 and LLM output = 1
- False negative (FN): evaluation label = 1 and LLM output = 0
- True negative (TN): evaluation label = 0 and LLM output = 0
- False positive (FP): evaluation label = 0 and LLM output = 1

We computed four standard metrics: accuracy, precision, recall, and the F1 score using the following formulas: accuracy = (TP + TN) / (TP + TN + FP + FN); precision = TP / (TP + FP); recall = TP / (TP + FN); and F1 score = 2 × (precision × recall) / (precision + recall).

In addition to the binary label, each model was prompted to output a probability value representing its self-reported confidence that the project was positive. Because this value was generated by the model itself, it should be interpreted as a self-reported score rather than a calibrated probability. Each model outputs the probability of the project being positive. We denote this probability by *p*. Values of *p* close to 0 indicate negative values and values close to 1 indicate positive values, whereas values close to 0.5 reflect low confidence in the model. To evaluate the relationship between the model’s self-reported confidence and classification performance, we computed the precision, recall, and F1 scores under two conditions for the same set of outputs. In the ALL condition, all 150 projects were included regardless of *p*. In the HIGH condition, projects with intermediate *p* values near 0.5 were excluded as ambiguous cases. Only projects satisfying *p* < 0.25 or *p* > 0.75 were retained in the high-confidence group, and projects with 0.25 ≤ *p* ≤ 0.75 were excluded from evaluation. These cut-offs were predefined practical cut-offs for confidence-based grouping and were not selected by data-driven threshold optimization. By comparing the F1 scores between the All and HIGH conditions, we quantitatively examined (i) whether models that produce many intermediate probabilities tend to show lower overall classification performance, and (ii) how much the F1 score improves when the evaluation is restricted to high-confidence predictions.

### Precision–recall curve and AUPRC analysis based on self-reported positive probabilities

To complement the evaluation of binary classification performance across all samples, we constructed precision–recall curves using each model’s self-reported positive probability as the prediction score. For each model, precision and recall were calculated across varying thresholds for the self-reported positive probability using the ground-truth labels of the projects, and the corresponding area under the precision–recall curve (AUPRC) was computed. We also generated model-specific precision–recall curves to evaluate whether the self-reported positive probability could function as a confidence indicator or ranking score in LLM-based metadata classification. The F1 scores compared with AUPRC were the conventional F1 scores calculated from the binary outputs for all 150 projects under prompt 2, rather than the F1 scores under the HIGH condition, in which samples with intermediate self-reported probabilities were excluded. The AUPRC values calculated here were not intended to replace the F1 scores computed from binary outputs across all samples, but were treated as complementary evaluation metrics based on the self-reported positive probabilities.

## Data availability

All supplementary data are available on Figshare (https://doi.org/10.6084/m9.figshare.30265717.v2).

Supplementary Figure 1: Self-reported probability and AUPRC analyses across the 17 evaluated models

Supplementary Figure 2: Inference runtime distributions and accuracy–runtime trade-off across models

Supplementary Table 1: Per-model counts of discrete self-reported probability values

Supplementary Table 2: Per-model AUPRC and F1/AUPRC ranking

Supplementary Table 3: Per-accession five-run reproducibility (n = 50)

Supplementary Table 4: Cross-session comparison between the original single-run outputs and the five-run reproducibility outputs

Supplementary Table 5: Detailed runtime statistics for open-weight models

Supplementary Table 6: Per-model accuracy and runtime summary (Prompt 2, ten locally executed open-weight conditions)

Supplementary Table 7: Example outputs of sample attribute extraction from metadata Supplementary Table 8: List of projects retrieved by keyword searches from the database Supplementary Table 9: Ground-truth labels of the benchmark dataset

Supplementary File 1: AUPRC analysis resources

Supplementary File 2: Reproducibility supplement (n = 50 reload-variance experiment)

Supplementary File 3: F1-vs-runtime analysis resources

Supplementary File 4: Integrated metadata text inputs used for LLM inference

Supplementary File 5: Prompts used for the classification and extraction tasks

Supplementary File 6: Workflow outputs and LLM-generated results for all model conditions

## Code availability

All codes in this study are publicly available at (https://github.com/mshintani22/open-weight-llm-metadata-curation-workflow) and are licensed under the MIT license.

## AUTHOR CONTRIBUTIONS

Conceptualization: M. S. and H. B.; methodology: M. S., D. A. and H. B.; software: M. S.; validation: M. S. and H. B.; formal analysis: M. S.; investigation: M. S.; resources: H. B.; data curation: M. S.; writing—original draft preparation: M. S.; writing—review and editing: M. S., D. A. and H. B.; visualization: M. S.; supervision: D. A. and H. B.; project administration: H. B.; funding acquisition: H. B. All authors have read and agreed to the published version of the manuscript.

## Acknowledgements

This work was supported by the Center of Innovation for Bio-Digital Transformation (BioDX), an open innovation platform for industry-academia co-creation (COI-NEXT); the Japan Science and Technology Agency (JST) [grant number JPMJPF2010]; and JST Broadening Opportunities for Outstanding young researchers and doctoral students in STrategic areas (BOOST) [grant number JPMJBS2424].

